# *Taf1* expression in mouse: novel transcripts and protein distribution

**DOI:** 10.1101/2025.09.08.674157

**Authors:** Peihang Li, Karen Cleverley, Elisa M. Crombie, Jan Bieschke, Elizabeth M.C. Fisher, Anna-Leigh Brown

**Affiliations:** Department of Neuromuscular Diseases and Queen Square Motor Neuron Disease Centre, UCL Queen Square Institute of Neurology, University College London, London WC1N 3BG, UK; LifeArc, Francis Crick Institute, 1 Midland Road, London NW1 1AT, UK; Institute of Prion Diseases, MRC Prion Unit at UCL, University College London, London W1W 7FF, UK

**Keywords:** TATA box binding-protein (TBP)-associated factor 1, TAF1, alternative splicing, truncated exon, alternative transcript

## Abstract

TATA-box binding protein associated factor 1 (TAF1) is the largest component of transcription factor IID (TFIID), a fundamental multiprotein complex for RNA polymerase II-mediated transcription. TAF1 is essential for promoter recognition, coactivator interaction, and normal development. Mutations in TAF1 cause developmental disorders and the lethal neurodegenerative disease X-linked dystonia-Parkinsonism (XDP). Our previous work suggested that this ∼170 kb gene has a more complex set of transcripts than currently catalogued. We therefore undertook a systematic assessment of *Taf1* transcription in mouse, given its widespread use as a model organism and its high genetic homology with humans. Using targeted nanopore sequencing, we reveal extensive transcriptional diversity and differential abundance of *Taf1* mRNAs across brain and body regions. We identify 19 novel transcript variants and multiple novel exons, and we predicted their protein domain architectures and modelled them using AlphaFold. Notably, we observed elevated RNA and protein expression in cerebellum compared to other brain regions. These findings substantially expand the transcriptional landscape of *Taf1* and provide crucial insights to guide investigation of transcript-specific mechanisms in neurodevelopmental and neurodegenerative disorders, enabling the creation of more accurate disease models.

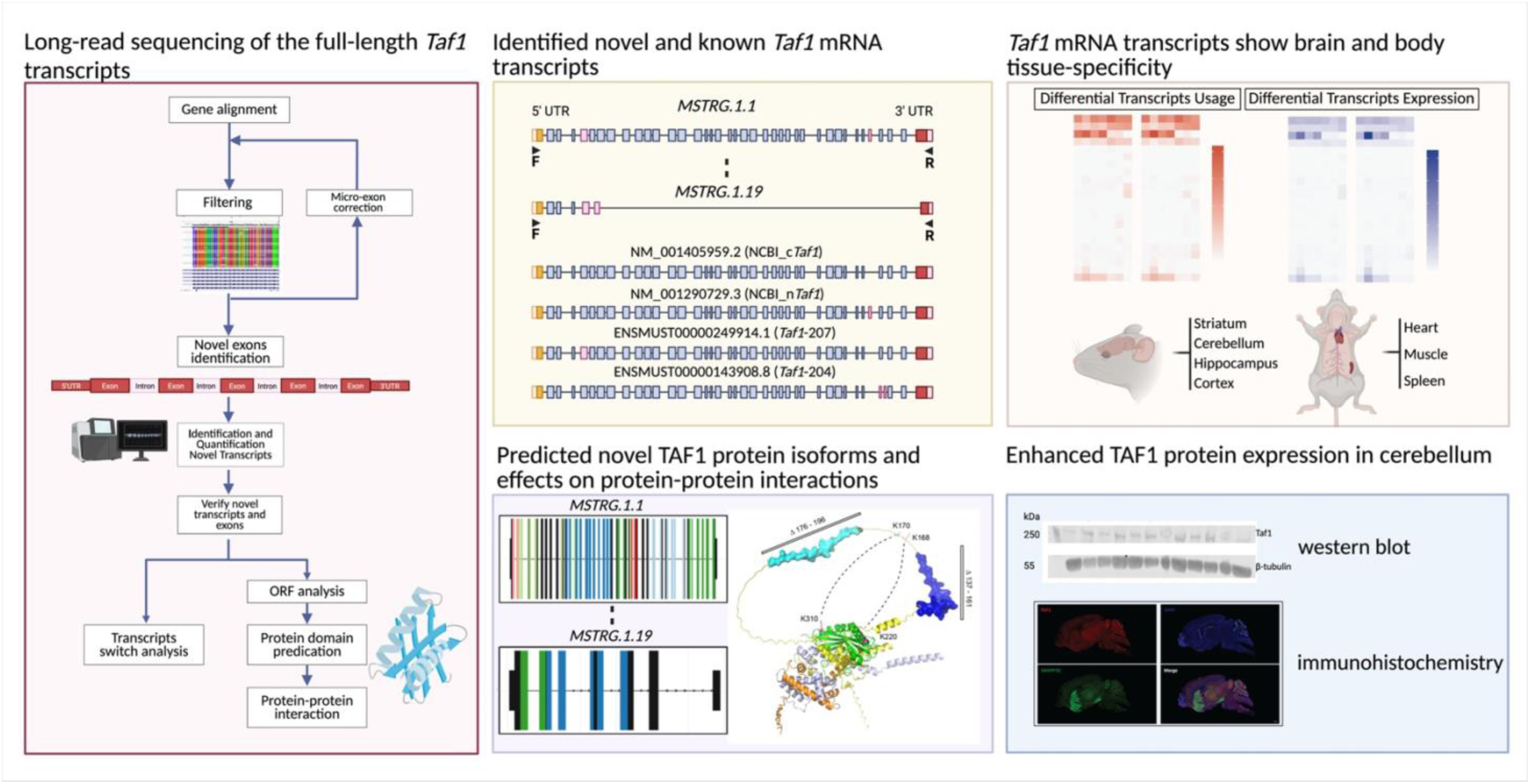

## Introduction

The TATA box binding-protein (TBP)-associated factor 1 gene (*TAF1*) is fundamental for the initiation of RNA polymerase II (pol II)-dependent transcription^1^. *TAF1* lies on the X chromosome and encodes a ∼250 kDa protein that is the largest constituent and key scaffold protein of the TFIID complex, a basal transcription factor for all polymerase II-mediated transcription in eukaryotes^2–4^. TAF1 has many functions involving protein-protein interactions^3,5,6^ and has DNA binding activity^4,7,8^. For example: the N-terminal domain of TAF1 binds TBP to inhibit DNA binding in its non-promoter bound state^9–11^; TAF1 has a specific TAF7-binding domain, and within this region a small winged-helix domain binds promoter DNA^12^; a zinc finger domain in TAF1 is involved in recruiting TFIID to endogenous promoters^13^. Furthermore, near the C-terminal of TAF1 lies a double bromodomain that recognises post-translational modifications – including acetylation, butyrylation, crotonylation – on histones and transcription factors^14–16^. Thus, through its diverse protein-domains, TAF1 plays an essential role in the regulation of transcription.

TAF1 is required for embryonic development^17–19^; female and male null mice die before E9.5^20^. Postnatally, gene editing and knockdown of *Taf1* in rat and mouse result in behavioural and neuronal deficits^21–23^. In humans, mutations in *TAF1* primarily affect males and missense and frameshift mutations cause the congenital disorder X-linked syndromic mental retardation−33 (MRXS33, X-linked intellectual disability, XLID), which presents with heterogeneous clinical features including facial dysmorphologies and cardiac defects^1,18,24–27^. In contrast, the adult-onset progressive neurodegenerative disorder XDP typically starts with dystonia (male mean age of onset 39 years of age) that generalises over 5-10 years then transitions into Parkinsonism, with a mean age of death of 56 years of age^28–30^. XDP arises from the insertion of a ∼2.6 kb SINE-VNTR-Alu (SVA) type F retroposon, into intron 32 of *TAF1*. This SVA carries between 30-55 CCCTCT repeats in its 5′ region and the number of these hexameric repeats is inversely correlated with the age of XDP onset^28,29^. The SVA insertion has a profound effect on transcription, producing aberrant mRNA variants; excision of the SVA in XDP iPSCs rescues *TAF1* expression, intron retention and aberrant splicing^28,31^. While it remains unknown how altered splicing leads to neurodegeneration, XDP is not thought to arise from a simple loss-of-function^28,29,32^.

Full knowledge of mRNA transcripts and protein isoforms within different cell types/life stages is essential for understanding gene function in health and disease. In writing a recent review of TAF1^1^ we realised that considerably more remains to be discovered about *TAF1* mRNA variants. The *TAF1* gene spans >166 kb on Xq13.1, and more than 38 exons have been annotated in NCBI GenBank/Ensembl. However, in humans at least, clearly more variants exist than have been described so far. For example, Capponi and colleagues amplified human prefrontal cortex cDNA between exons 30 to 38 only and sequenced the ten most abundant products, finding previously unreported exons and transcripts^33^.

Alternative splicing (AS) is a feature of 95% of human pre-mRNAs from multi-exon genes, resulting in variants with different UTRs, retained introns, skiptic exons, cryptic exons, mutually exclusive exons, and alternative polyadenylation sites. This process leads to immense transcriptional complexity for each cell type^34^, with highest splicing diversity seen in the central nervous system^35,36^, perhaps reflecting the massive variety of cell types in this tissue^37^. Humans average >8 mRNA variants per protein coding gene^38^; AS is highly species specific and this average is likely lower in mice^37,39^. To comprehend the biology of each gene, it is essential to understand the role each mRNA transcript/protein isoform plays in an individual cell type, noting that proteins from a single gene can play opposing roles to control specific pathways. For example, depending on the mRNA transcript, the *BCL2L1* gene can encode either a pro- or anti-apoptotic protein isoform^40^. Furthermore, individual protein isoforms have their own independent interactomes^38^. Thus, we need a systematic classification of *TAF1* protein-coding transcripts, spatially and temporally.

With the advent of long-read sequencing it is becoming easier – although still not straightforward – to define the transcriptional complexity of a single gene^41^. Challenges include the high error rates of sequencing reads, and, for example, the difficulty in detecting microexons (exons <30 bp)^36^ within mRNAs/cDNAs of several kb. Using the mouse, a key model for human disease, we catalogued *Taf1* transcripts across male and female brain and non-brain tissues from young adults. By amplifying long cDNAs between the *Taf1* 5′ and 3′ UTRs and sequencing them with nanopore technology, we developed a bioinformatics pipeline for analysing novel transcripts and predicting protein structures, discovered new exons and mRNA transcripts, evaluated their expression patterns across tissues, and identified inconsistencies in existing annotations that warrant further experimental validation. This work builds on our previous description of mouse *Taf1* transcripts^1^, adding semi-quantitative data, including the unexpected finding of particularly high transcript and protein abundance in the cerebellum. Together, these results contribute to a more complete understanding of TAF1 and highlight the complexity of this essential gene/protein that has multiple functions.

## Results

### Generation of mouse *Taf1* cDNAs from brain and non-brain tissues and long-read nanopore sequencing

While the *TAF1* gene is ubiquitously expressed, *TAF1* mutations show selective tissue pathology; notably, XDP is reported to primarily affect the striatum, and congenital *TAF1* mutations affect the heart and other tissues. Therefore, to assess tissue-specific splicing patterns, we collected RNA from multiple brain (striatum, cerebellum, hippocampus, cortex) and body (heart, spleen, quadricep muscle) regions of 3-month-old (young adult) male and female C57BL/6J mice, n=2 per sex (Supplementary Table S1). We created cDNA by reverse transcribing from oligo-dT primers, then amplified the product between PCR primers that bound to the 5′ UTR and 3′ UTR at chrX:100.576.386 and chrX:100.643.185, respectively (Figure 1A). Amplification between these primers gives a product of 5,856 bp based on GenBank *Taf1* ‘mouse canonical transcript’ (NM_001405959.2). Long-read nanopore sequencing generated a total of 2 million reads, with an average of 72,411 reads per tissue and average read length of 4,700 bp, mean read quality was 20 – representing a base call accuracy of 99% (Supplementary Table S2)

**Figure 1.**
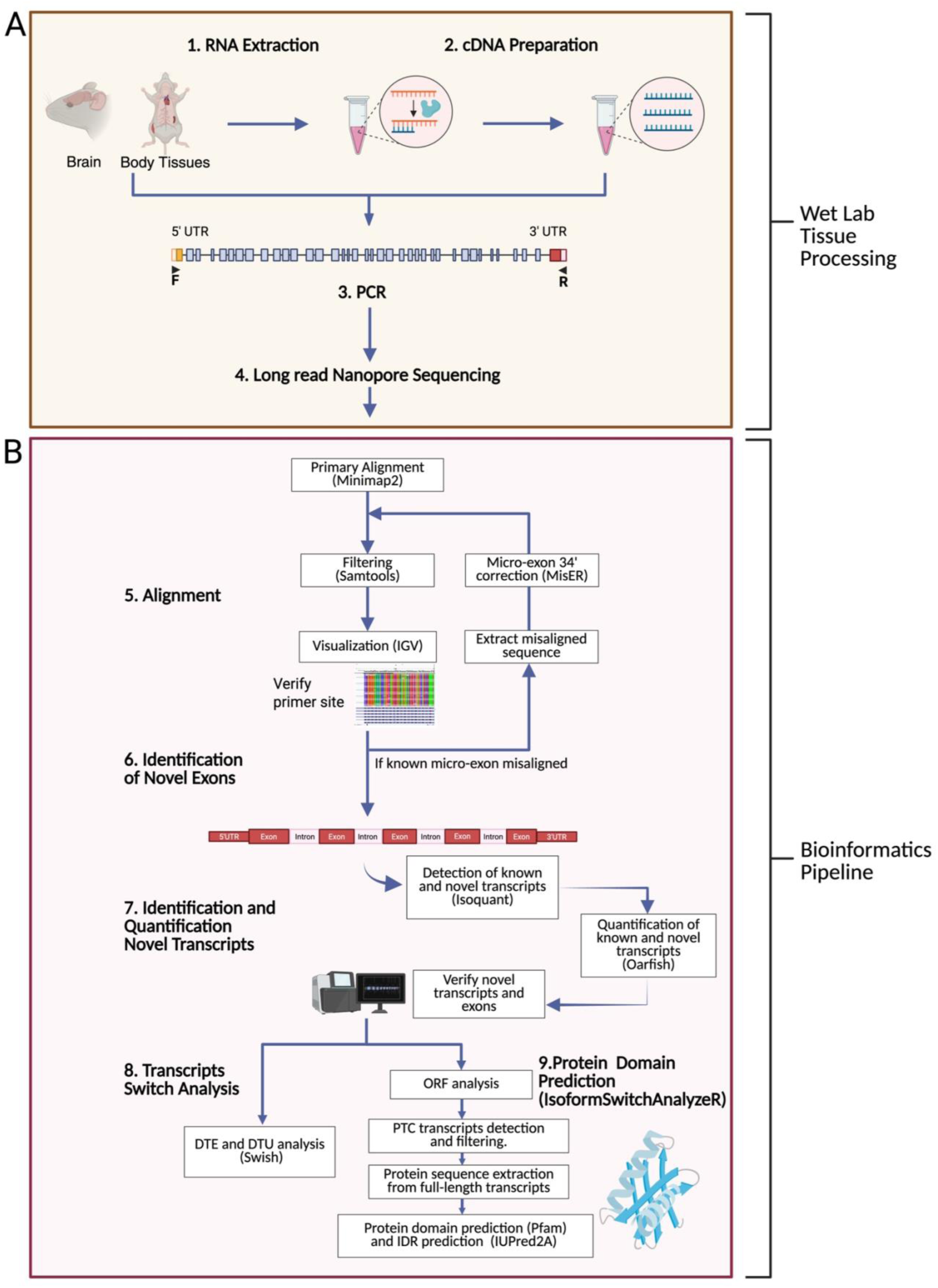
Sequencing strategy and bioinformatics pipeline for identifying and quantifying *Taf1* transcripts using long-read sequencing. (**A**) RNA was extracted from brain and body tissues and used for cDNA synthesis from oligo-dT primers. cDNA was amplified by PCR using primers designed to the 5′ and 3′ UTRs based on mouse genome assembly GRCm39. (**B**) Long-read nanopore sequencing of PCR products generated data that were aligned to GRCm39 via minimap2. Alignments were filtered with samtools and visualized in IGV to verify primer specificity and exon-intron structures. Misaligned reads, particularly those indicating the potential presence of microexon 34′ (6 bp), were corrected using MisER. Transcript analysis was conducted using Swish to assess differential transcript usage (DTU) and differential transcript expression (DTE) across specific tissue regions. Protein domain prediction was performed through IsoformSwitchAnalyzeR, integrating open reading frame (ORF) predictions and PFAM domain annotations. Protein structure prediction employed IUPred2A. PTC: premature termination codon.

### Targeted long-read analyses reveal novel *Taf1* transcripts in brain and non-brain tissues

To comprehensively analyse *Taf1* transcript expression, long-read sequencing data were aligned to GRCm39 using minimap2. Given the challenge of aligning microexons with minimap2, noting the 6 bp microexon 34′ within *Taf1*, MiSER was incorporated to improve mapping accuracy for microexons. Following alignment, IsoQuant was employed for transcript assembly and then assembled transcripts across samples were merged using StringTie2. Transcripts that appeared in multiple samples were kept for downstream analyses and expression quantification. After assembling a single set of known and novel *Taf1* transcripts, Oarfish was used for transcript quantification and Swish for DTU and DTE. Further functional interpretation, including ORF and protein domain prediction, was performed using IsoformSwitchAnalyzeR (Figure 1B). This integrated pipeline enabled robust identification and quantification of annotated and novel *Taf1* transcripts, tissue specific expression, and prediction of their protein-coding consequences and domain architectures.

Ensembl currently shows three full-length *Taf1* mRNA variants (*Taf1*-201, *Taf1*-205, *Taf1*-207), while GenBank lists five (Variants 1-5)^1^. These variants primarily differ in whether: exon 5 is full-length or shortened by 63 bp at its 5′ end; the presence of the 6 bp microexon, 34′, which is observed exclusively in the neuronal form of *Taf1* (n-*Taf1*); an interruption within exon 38, noted only in GenBank Variants 4 and 5. This exon 38 interruption likely results from an amino acid sequencing or alignment artefact, as it lacks support from genomic DNA sequencing.

Curiously, it is not clear where the *TAF1*/*Taf1* ORF begins. For example, Ensembl annotates two human mRNA variants, *TAF1*-204 (c*TAF1*, NM_004606.5, GenBank ‘Variant 1’) and *TAF1*-228, which have identical DNA sequences. However, *TAF1*-228 possesses a predicted start codon (ATG, methionine) 60 bp (20 amino acids) upstream of the predicted start ATG in *TAF1*-204. Currently, there is no experimental evidence from human, mouse, or other species to confirm which of these potential start ATGs is used *in vivo*. The downstream ATG (as found in *TAF1*-204) is associated with a more favourable Kozak sequence^42,43^, which typically enhances translational efficiency. Mouse *Taf1* transcripts similarly contain two potential N-terminal start codons, separated by 60 bp (20 amino acids), and it remains unclear if one or both are active. Our 5′ primer was designed to terminate 19 nucleotides upstream of the first annotated start codon, and so all novel transcript sequences identified in this study encompass both potential start codons.

Across the tissues we analysed, four known *Taf1* transcripts were expressed in at least two tissue types, three maintaining the ORF (GenBank reference ‘canonical’ transcript *cTaf1*, *nTaf1, Taf1-207*) and one *(Taf1-20*4, Figure 2) is predicted to result in nonsense-mediated decay (NMD).

**Figure 2.**
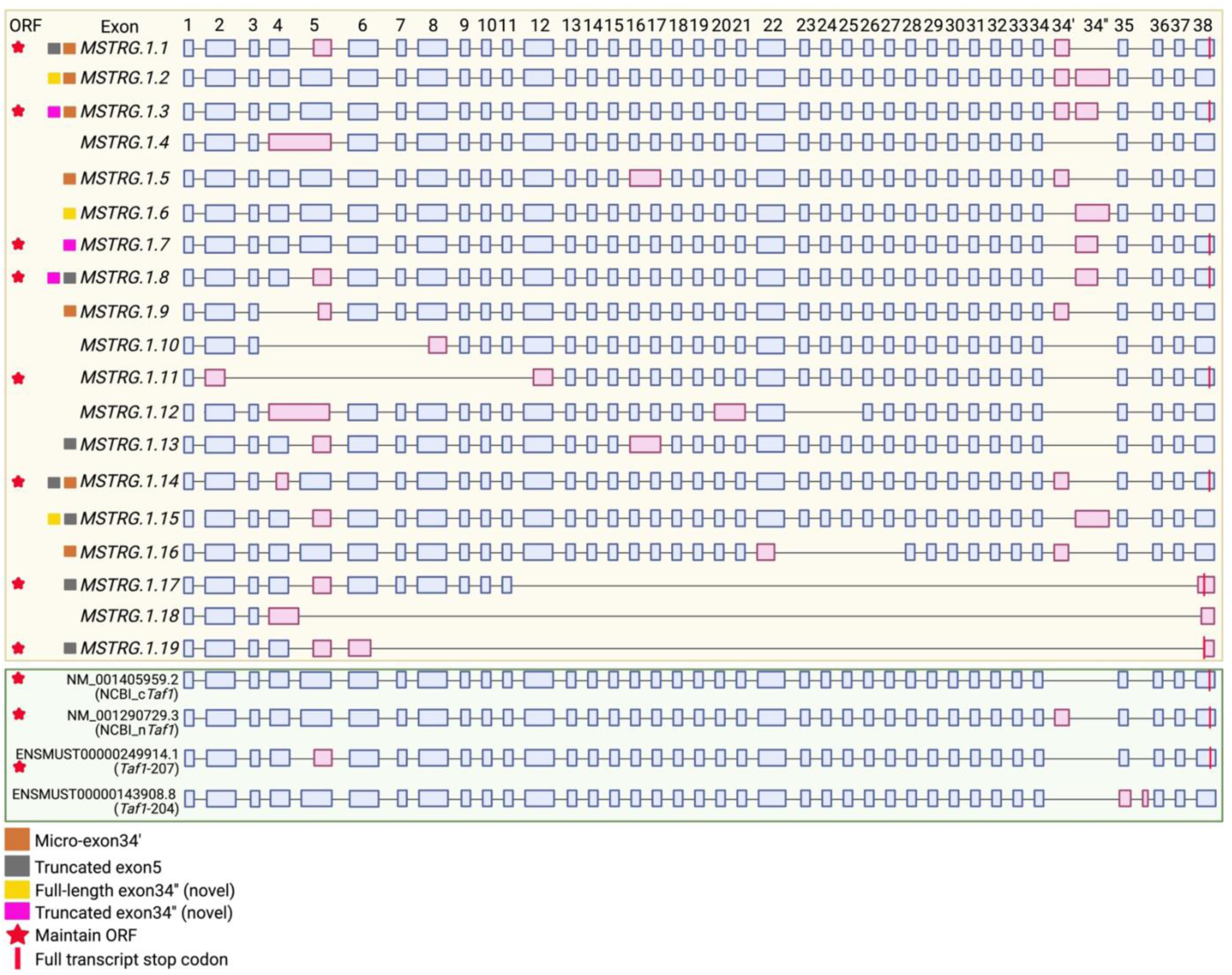
Intron-exon structures of novel and known *Taf1* transcripts. The top panel (yellow) depicts 19 novel *Taf1* transcripts identified through our analysis of mouse brain and body regions. Exons are represented as boxes, with exons differing from the canonical GenBank reference transcript (c*Taf1*, NM_001045959.2) highlighted in red. Alternative exons repeatedly identified across multiple transcripts are color-coded by the boxes at the start of the line: truncated exon 5 (grey), neuron-specific microexon 34′ (brown), novel long exon 34′′ (yellow), and novel short exon 34′′ (purple). Variants marked with a red star maintain an intact ORF without premature termination codons (PTCs), indicating potential functional, full-length transcripts. Red vertical bars indicate positions of stop codons at the end of the ORFs in the full-length transcripts. The lower panel (green) shows four known *Taf1* transcripts in GenBank/Ensembl databases. Transcript n*Taf1* (NM_001290792.3) incorporates microexon 34′, *Taf1*-207 (ENSMUST00000209949.1) contains truncated exon 5, and *Taf1*-204 (ENSMUST00000143908.8) is likely subject to NMD. ‘*MSTRG*’ is a transcript identifier assigned by StringTie software during transcript assembly. Supplementary Table S5 shows the length of the novel exons.

Beyond the known transcripts in Ensembl/GenBank, we identified 19 novel *Taf1* transcripts (Figure 2). Six of these novel transcripts (*MSTRG.1.1, 1.3, 1.7, 1.8, 1.11, 1.14*) contain an ORF extending from exon 1 to exon 38. Exon 38 is the most 3′ exon identified in our analysis and includes the 3′ UTR. *MSTRG.1.1, 1.3, 1.7, 1.8,* and *1.14* comprise 39 or 40 exons, resulting in predicted protein translations between 1,890 and 1,952 amino acids. These translations exhibit high sequence similarity (96.1% to 97.7%) to the canonical cTAF1 protein. In contrast, *MSTRG.1.11* features an ORF from exon 1 to exon 38 but skips exons 3 through 11, leading to a shorter predicted translation of 1,358 amino acids.

Two novel transcripts (*MSTRG.1.17, MSTRG.1.19*) have markedly few internal exons and an upstream stop codon in a truncated exon 38 giving rise to a shorter ORF. While these transcripts use an upstream stop codon from *cTaf1*, they are not predicted to undergo NMD. Percentage similarities of *MSTRG.1.11, 1.17* and *1.19* to *cTaf1* are shown in Supplementary Table S4.

Alternative exons incorporated into the new eight likely coding variants include truncated versions of exons 4, 5, 6, microexon 34′ (6 bp) and a long or short form of a new exon, 34′′. In addition to skipping internal exons, transcript *MSTRG.1.11* has a 3′ truncation of exon 2 and a 5′ truncation of exon 12.

Thus, we have identified both short (42 bp) and long (120 bp) forms of exon 4, short (179 bp) and long forms (242 bp) of exon 5 – all of which maintain the ORF, and found a new exon, 34’′, which also has short (105 bp) and long (172bp) forms but only the short form maintains the ORF. Based on their sequences, novel mRNA transcripts *MSTRG.1.4, MSTRG.1.5, MSTRG.1.12* and *MSTRG.1.13* are predicted to undergo NMD.

### Mouse *Taf1* splicing transcripts and abundance vary across tissues

To investigate the tissue-specific expression pattern of *Taf1* transcripts, we calculated DTU and DTE using the R package Swish. DTU reflects shifts in transcript proportions within a region, whereas DTE indicates differences in the absolute expression levels of individual transcripts^44^. In Figure 3 we show *Taf1* transcript abundance in seven tissues in female and male mice. We went on to compare DTU and DTE between specific tissues, for example, cerebellum values versus the three other brain regions as well as with combined body/brain regions; only significant differential expression is shown (Figure 4). In this analysis we did not find significant sex differences.

**Figure 3.**
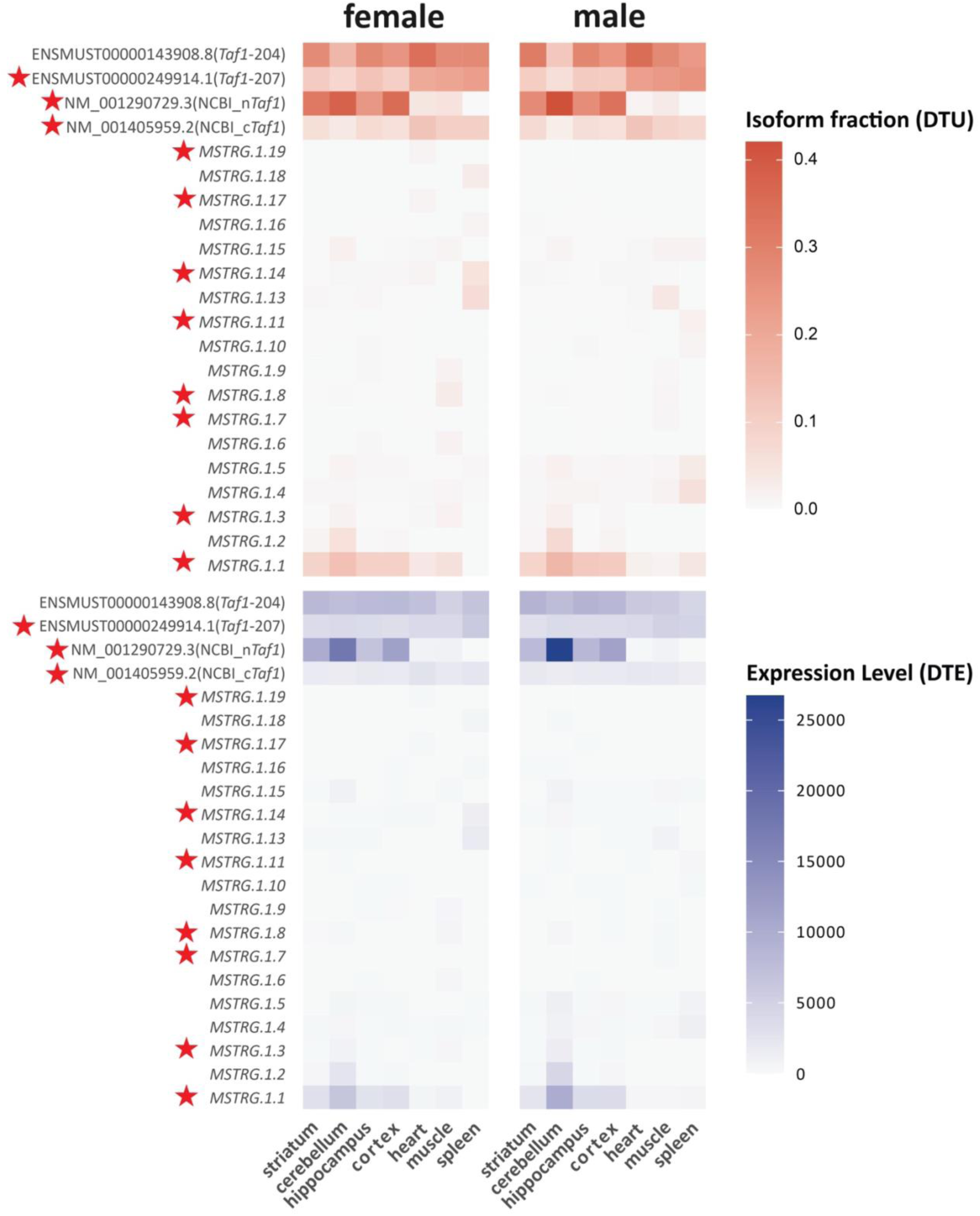
DTU and DTE for *Taf1*. The top DTU panel illustrates transcript abundance (red gradient), quantified from normalized Oxford Nanopore Technology (ONT) transcript reads, for both annotated and novel *Taf1* transcripts across seven distinct tissues (striatum, cerebellum, hippocampus, cortex, heart, muscle, and spleen) in male and female mice. The bottom DTE panel presents transcript fractions (blue gradient), indicating the proportional representation of each transcript within these tissues. Red stars denote transcripts that maintain ORFs.

**Figure 4.**
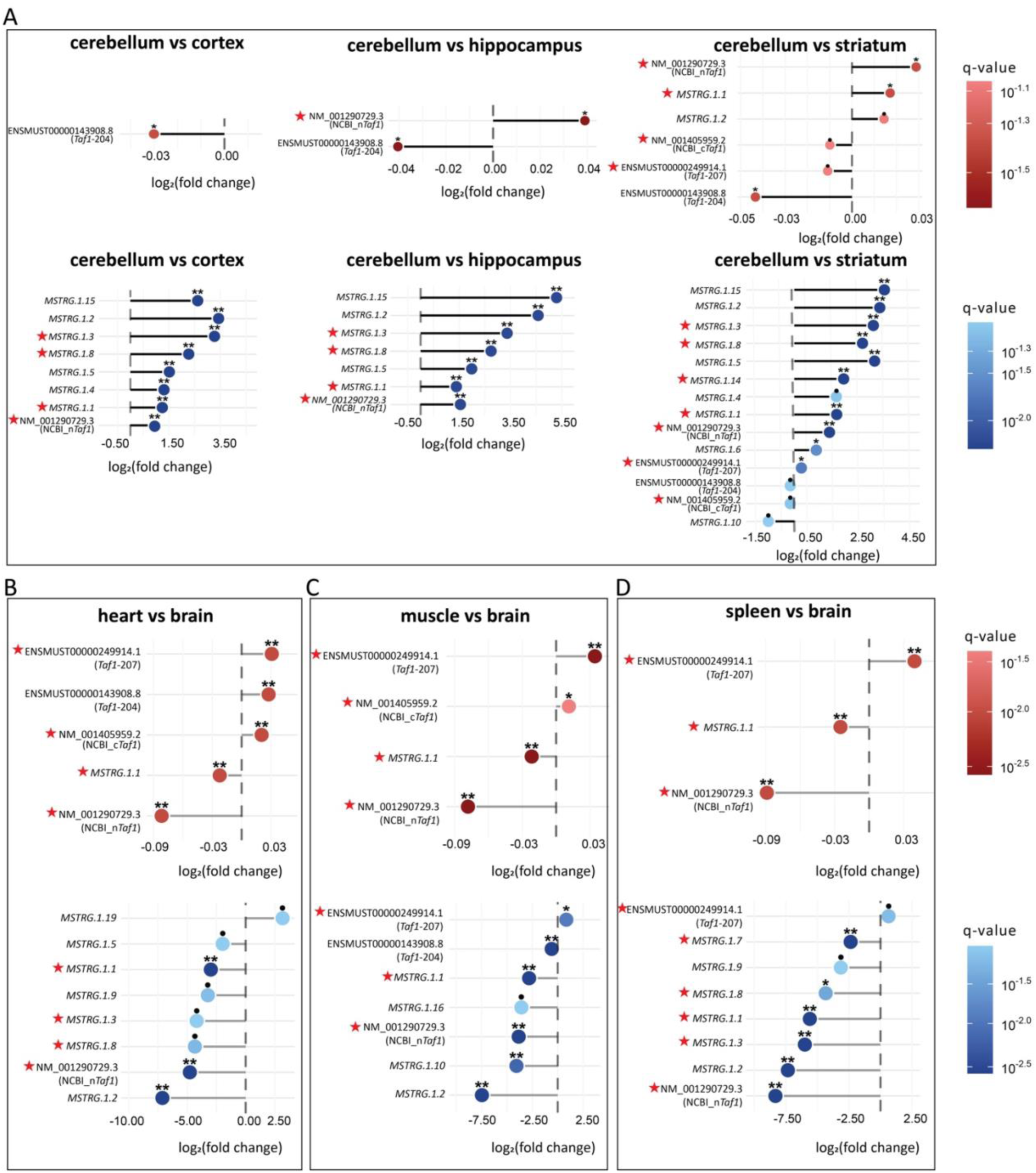
Lollipop plots of significant DTU (red) and DTE (blue) data across selected tissue comparisons. (**A**) Transcript-level comparisons between the cerebellum and brain regions (cortex, hippocampus, and striatum). (**B**) Heart tissue compared with brain regions. (**C**) Muscle tissue compared with brain regions. (**D**) Spleen tissue compared with brain regions. Log2 fold changes are presented along with corresponding q-values, indicated by colour gradient intensity. Asterisks indicate significance: **q < 0.01, *q < 0.05, •q < 0.1. Full length real transcripts (red star).

Surprisingly, we found that cerebellum had the highest overall expression of *Taf1* mRNA as well as transcript diversity in the brain regions (9 out of 19 novel transcripts detected had expression levels of 0.5 to 5.5 DTE fold change, therefore we chose to use this region as the baseline tissue for additional transcript analyses).

For the four known *Taf1* transcripts, we confirmed that *nTaf1* (NM_001290792.3) is neuronally enriched (Figure 3A; B) and furthermore, is enriched in the cerebellum compared to the other brain regions, as shown by significant DTU differences in hippocampus and striatum (q-value < 0.05) and significant DTE differences in cortex, hippocampus, and striatum (q-value < 0.01) (Figure 4A). Both *Taf1-207* and *cTaf1*, lacking exon 34′, are the major protein-coding transcripts in body regions (Figure 3; Figure 4). However, *Taf1-207,* which has a truncated exon 5, not *cTaf1*, is the most abundant protein coding variant in the body regions (Figure 3; Figure 4). The known NMD-inducing transcript, *Taf1-204,* shows lower expression in the cerebellum with significantly higher usage observed in the striatum (DTU, q-value < 0.05; Figure 4A).

Among the 19 novel *Taf1* transcripts, *MSTRG.1.1* and *MSTRG.1.2* have brain-specific expression when compared to heart, muscle, and spleen, with highly significant DTU and DTE values in brain compared to peripheral body regions (q-value < 0.01; Figure 4B-D). Of the novel transcripts, *MSTRG.1.1* has the highest expression across most tissues (Figure 3). Six novel transcripts (*MSTRG.1.1*, *1.2*, *1.3*, *1.5*, *1.8*, *1.15*) have markedly high expression in the cerebellum relative to the other brain regions, showing log2 fold changes ranging from ∼1.5 to >5.5, all with highly significant DTE (q-values < 0.01). Three of these six transcripts (*MSTRG.1.1*, *MSTRG.1.3*, and *MSTRG.1.8*) are full-length and predicted to encode functional TAF1 proteins.

Intriguingly we find microexon 34′ is present in eight transcripts: *nTaf1*, *MSTRG.1.1, 1.2, 1.3, 1.5, 1.9, 1.14, 1.16* only four of which likely produce full-length proteins (*nTaf1*, *MSTRG.1.1, 1.3, 1.14*). These transcripts tend to be expressed in brain rather than body tissues (Figure 3; Figure 4).

### Mouse TAF1 protein expression is higher in cerebellum

To assess TAF1 protein distribution in mouse brain, we performed western blotting with antibody TAF1 [177-4] raised to the epitope TPGPYTPQPPDLY, derived from peptides encoded by exons 34 and 35 of *TAF1*/*Taf1* (*Taf1*-201 and *Taf1*-202)^45^ (Figure 5A; Supplementary Table S3). At the resolution of our gels, this antibody detected a single TAF1 band of ∼250 kDa (we could not separate proteins with/without the possible 20 amino acids at the carboxy terminus). Reproducible results were obtained through multiple blots; a representative blot of 3-month-old male tissue is shown with quantification in Figure 5 (n=3). We analysed data from 6 males and 6 females, at 3-, 6-, and 12-months old, to assess changes over lifespan – noting our limited ability to detect different isoforms (Supplementary Figure S1A; S1B).

**Figure 5.**
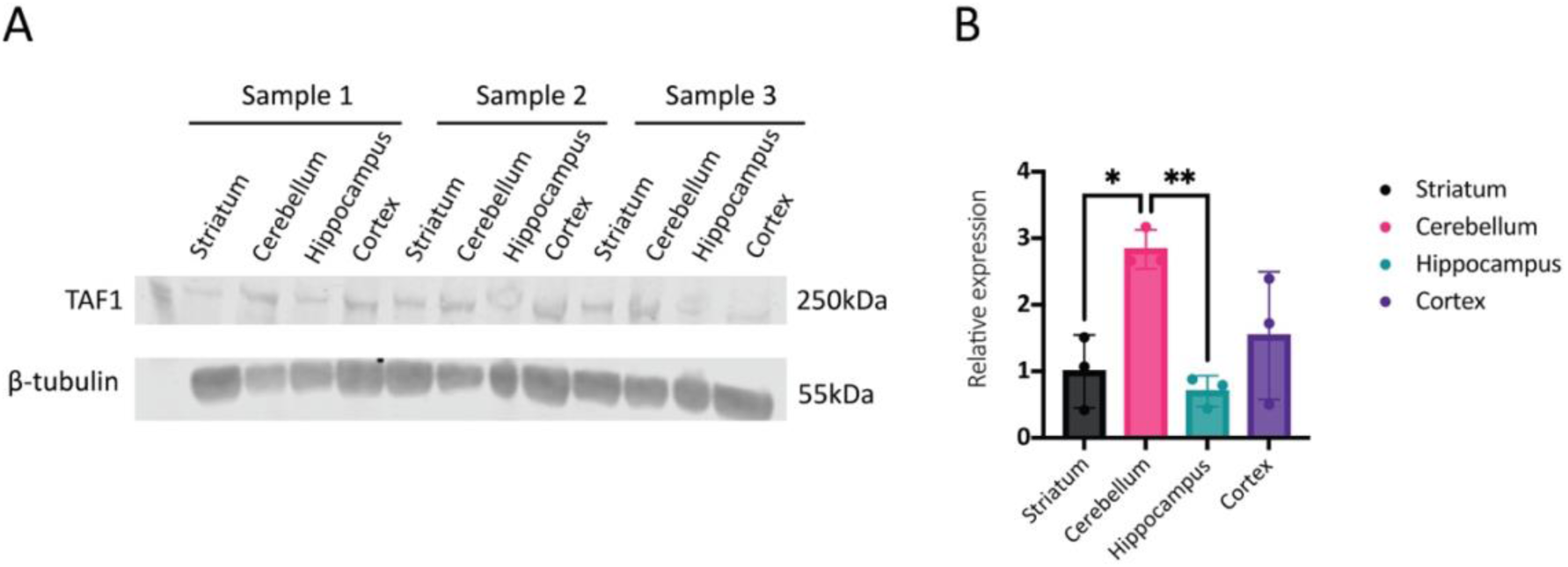
Differential Expression of TAF1 in brain regions of 3-month-old male mice. (**A**) Detection of TAF1 protein expression in striatum, cerebellum, hippocampus and cortex by western blot (n=3). Representative blot with β-tubulin used as a loading control. (**B**) Quantification of western blots (n=3), bar graphs represent the mean ± standard deviation (SD) of the relative expression levels. The data for each brain region are: striatum (black), cerebellum (pink), hippocampus (blue), and cortex (purple). Significant differences between brain regions are indicated by asterisks, with statistical significance denoted as \**p* < 0.05, \*\**p* < 0.01.

In line with our mRNA results, cerebellum consistently exhibited the highest levels of TAF1 protein across all ages and both sexes: expression was significantly elevated in 3-month-old males (Figure 5A; 5B) and 6-month-old male and female groups (Supplementary Figure S1A; S1B) and peaked at almost three times the relative level compared to striatum. Cerebellum showed the most statistically significant differences compared to the other brain regions, particularly cerebellum and cortex, where the difference reached the highest statistical significance in 6-month-old males [F(3,8) 27 = 19.40, *p* = 0.0005] and 6-month-old females [F(3,7) = 5.20, *p* = 0.0324] (Supplementary Figure S1A; S1B). TAF1 expression in the cerebellum was also markedly higher compared to striatum and hippocampus, particularly in 3-month-old males (Figure 5A; 5B), where it was 3.6 times higher than in the striatum [F(3,8) = 7.80, *p* = 0.0203] and 3.8 times higher than in the hippocampus [F(3,8) = 7.80, *p* = 0.0089]. Interestingly, in 12-month-old males and females, the difference between the cerebellum and cortex was less pronounced (Supplementary Figure S1A; S1B).

The hippocampus also had relatively high TAF1 expression, especially in male mature adult mice (6 months of age). The difference between hippocampus and cortex was statistically significant, with TAF1 expression in hippocampus being ∼1.5 times higher than in cortex [F(3,8) = 19.40, *p* = 0.0014] (Supplementary Figure S1A; S1B). However, in older adult mice (12 months), hippocampal TAF1 expression was much lower than in cerebellum and cortex, which contrasts with its expression trend in younger and mature mice. In contrast, the cortex and striatum showed relatively lower levels of TAF1 expression compared to cerebellum (less than 2-3 times lower) and hippocampus (less than 0.5-2 times lower), except in 12-month-old male mice. Interestingly, the striatum generally exhibited slightly higher TAF1 expression than the cortex, in all age groups of females and with statistical significance in 6-month-old males [F(3,8) = 19.40, *p* = 0.0055] (Supplementary Figure S1A; S1B).

To validate the significant relative TAF1 expression observed in the cerebellum compared to other brain regions, immunohistochemistry was conducted on the left hemisphere of 3-month-old male and female C57BL/6J mice (n=3/sex) (Supplementary Figure S2); sagittal sections allowed the cerebellum to be visualized with other brain regions. DAPI staining identified nuclei, with immunofluorescent staining for TAF1 and DARPP-32 [H-3] – a marker of medium spiny neurons in the striatum (Supplementary Table S3). This analysis verified that nuclear TAF1 protein expression was elevated in the cerebellum compared to hippocampus and striatum, as indicated by co-localization of DAPI with TAF1 (Supplementary Figure S2A; S2B). We detected no significant differences between males and females. These immunohistochemistry findings align with the western blot data, confirming that TAF1 is highly expressed in the cerebellum, relative to other brain regions.

### Novel TAF1 isoforms may have different protein interactions

We conducted protein domain prediction analysis for the novel transcripts predicted to maintain ORFs (*MSTRG.1.1, MSTRG.1.3, MSTRG.1.7, MSTRG.1.8, MSTRG.1.11, MSTRG.1.14, MSTRG.1.17, MSTRG.1.19*). Using PFAM domain analysis, these novel *Taf1* transcripts – except *MSTRG.1.17* and *MSTRG.1.19* – share conserved features with the canonical GenBank *cTAF1* transcript (NM_001045959.2): two bromodomains; one TBP-binding domain; one zf-CCHC_6 zinc finger domain. The variations between *MSTRG.1.1* and *MSTRG.1.14* were primarily in the intrinsic disordered regions (IDRs) and IDRs with binding regions (Figure 6A). Both *MSTRG.1.17* and *MSTRG.1.19* lack bromodomains, DUF3591, and zinc finger domains, with *MSTRG.1.19* additionally lacking an IDR binding region.

**Figure 6.**
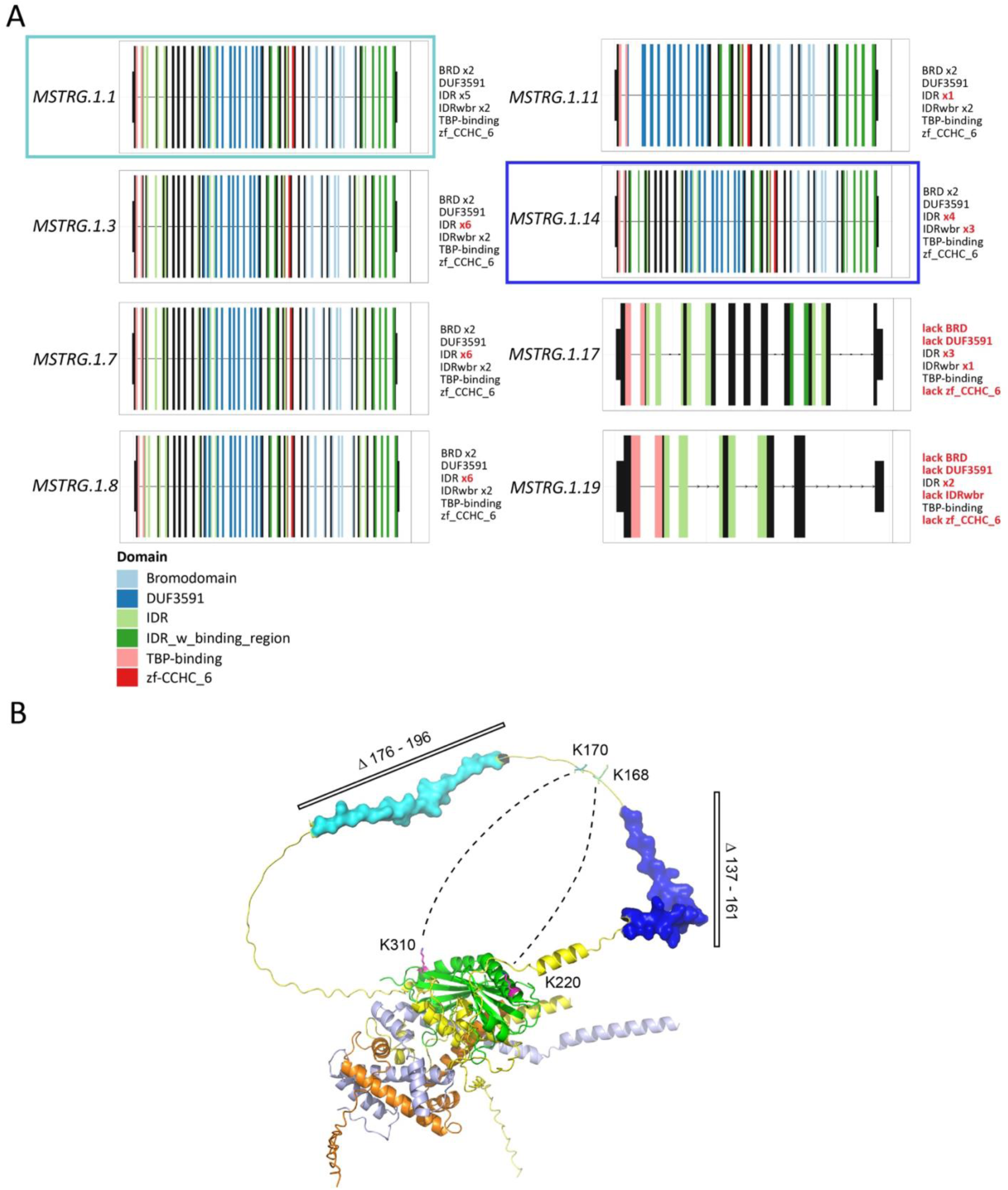
TAF1 predicted protein domains. (**A**) Predicted protein domains (using PFAM) for predicted ORFs within novel *Taf1* transcripts (Bromodomains; Domain of Unknown Function, DUF3591; Intrinsic Disordered Region, IDR; Intrinsically Disordered Region with a binding region, IDRwdr; Tata-Box Protein binding domain, TBP; zinc finger domain, zf-CCHC_6). *MSTRG.1.1* (light blue box) and *MSTRG.1.4* (blue box) are highlighted, which were used for protein–protein interaction analysis shown in panel (B). (**B**) Complex of mouse TAF1 (amino acids 1-300, yellow), TBP (amino acids 141-320, green), TAF11 (amino acids 50-211, grey) and TAF13 (full length, orange) modelled in Alphafold 3. Regions deleted in TAF1.1 (light blue) and TAF1.14 (blue) both lie in an unstructured loop between the TAF1 N-terminal domain (TAND) and the putative interaction motif of TAF1 with TAF11 and TAF13. Dashed lines indicate putative interactions between TAF1 and TBP derived from chemical crosslinking^3^.

We next sought to predict if any of these protein isoforms might affect TAF1 protein-protein interactions. Most predicted TAF1 isoforms varied in the C-terminal IDRs, which are notoriously difficult to model, and for which interaction partners are often unknown. We therefore focused on the variation in the N-terminus region of TAF1, for which protein structures of the putative protein complex between TAF1, TAF11, TAF13 and TBP are available^3, 4^. Amongst these, known transcript *Taf1-207* and novel transcript *MSTRG.1.1* contain a truncated exon 5 and *MSTRG.1.14* contains a truncated exon 4. These two exons code for amino acids that lie in the intrinsically disordered loop region between TAND and TAF11/13 interacting domains of TAF1 and would shorten the loop by either 23 amino acids (truncated exon 4, *MSTRG.1.14*) or 20 amino acids (truncated exon 5, *Taf1-207*; *MSTRG.1.1*; Figure 6B) ^47^.

## Discussion

RNA splicing allows a single gene to produce numerous RNA transcripts and protein isoforms which can have profound effects on a protein’s interaction partners, localisation, and function. TAF1 is an essential protein for healthy development, and results in wide-ranging phenotypes when mutated. Little is known of its transcripts and protein isoforms, or its regulation. To characterise *Taf1* gene expression in male and female adult mice we created long mRNA, reverse transcribed long cDNA from oligo-dT primers, amplified cDNA between 5′ and 3′ UTRs and then used nanopore sequencing to assess transcripts in mouse brain and non-brain tissues chosen for their relevance to *TAF1* disease in humans. This provides us with a snapshot of the complexity of *Taf1* expression, given that we are looking at one age (3-months old) and could not detect transcripts with alternative 5′ or 3′ UTRs or polyadenylation sites. Nevertheless, we have now identified 23 transcripts in mouse, of which only four are described in GenBank/Ensembl.

*Taf1* splicing presented a bioinformatic challenge because transcripts are predicted to be up to 5.9 kb and can include the neuronally-enriched microexon 34′ which is only 6 bp in length. Microexons shorter than 20 nucleotides are especially prone to being missed in standard long-read sequencing analyses without dedicated correction strategies^46^. In our analyses of *Taf1* diversity, we aimed to both identify and accurately quantify differential transcript usage across tissues. Thus, we implemented a strategy that combined reference-guided transcript assembly with transcript-level quantification, using a tool specifically optimized for long-read data. This approach allowed us to capture subtle but biologically meaningful differences in transcript expression that might be missed by methods focusing solely on annotation or quantification alone.

Several novel transcripts were identified with low expression levels, consistent with the general trend that novel mRNA variants exhibit lower expression compared to annotated transcripts^47–49^. We identified 19 novel mRNAs, of which eight are predicted to encode ORFs, two of these encode notably short proteins, which only contain the TBP-binding and IDR domains of the TAF1 protein. The other novel transcripts are predicted to undergo NMD and therefore are likely important for controlling TAF1 protein levels^50^. Of the four known variants, three maintain a full-length ORF and one, *Taf1*-204, has a PTC. Overall, we have identified short and long forms of known and novel exons and found typical AS events such as intron skipping/retention, that indicate considerable complexity for *Taf1* regulation and function.

Our expression analysis of the 23 transcripts in 3-month-old mice showed notable differences in their tissue distribution, with some largely brain specific (for example, *MSTRG.1.1* and *nTaf1 -*NM_001290792.3) whereas others appear present in all tissues (for example, *Taf1*-204). The four mRNA variants previously described tended to be more abundant than the 19 novel variants. Overall, in the brain tissues, cerebellum was the site of highest *Taf1* expression. This was a surprise as data from human brains affected by XDP indicate that striatum is the key site of pathology^51–53^. Intriguingly, the NMD-associated *Taf1*-204 transcript showed highest abundance in the striatum relative to cerebellum, perhaps indicating a regional specific vulnerability for loss of function mutations as a potential pathogenic mechanism in XDP. Heart is consistently affected in congenital cases of human *TAF1* mutation, and we note the relatively high expression of *Taf1*-204 in this tissue as well.

We analysed overall protein expression in mouse brain, and though currently available TAF1 antibodies do not allow us to distinguish protein isoforms, we determined that cerebellum showed consistently higher expression of TAF1 than other brain regions. This increased protein abundance was consistent across ages and sex in 3-, 6- and 12-month-old male and female mice. Some variation was apparent in expression levels in cortex, striatum, and hippocampus, but immunohistochemistry of male and female mouse brain at 3-months of age supported higher TAF1 expression in the cerebellum. Our results do not allow us to address cell type for AS, but such events and the resulting protein isoforms may be entirely cell-specific and related to key cellular functions^54^.

We aligned *Taf1* transcripts with known protein domains and determined that in novel transcripts *MSTRG.1.1* and *MSTRG.1.14* truncations in exons 4 and 5 both shorten the loop connecting TAND and TAF11/13 interacting domains of TAF1, possibly stopping interaction with TBP, TAF11 or TAF3^3^. There may be other interaction partners in the full TFIID complex, which could be affected by these deletions^4^. The shortened loop may reduce the conformational flexibility of TAF1 within TFIID and impair tethering of TBP by TAF1 in the absence of TAF11/13 binding. Furthermore, the deletions could interfere with a putative interaction between K168/K170 of TAF1 and K220/K310 (K243/K333 in human TBP)^3^, potentially altering TFIID function.

In summary, we find considerably more complexity in transcription from the mouse *Taf1* locus than previously reported. This includes transcripts that have full-length ORFs but different domain composition and those that likely regulate TAF1 levels through NMD. We confirm that microexon 34′, which lies within an IDR and is also present in human *TAF1*, is present within approximately a third of our transcripts including *nTaf1* and seven other transcripts. Of these eight transcripts, seven show significantly elevated expression in brain regions/nervous system-specific compared to body regions. The inclusion of microexons in mature mRNAs is frequent in the CNS and the encoded residues are often located on protein surfaces to regulate protein-protein interactions^36^.

We note that many of the protein-coding predicted protein isoforms varied primarily in the length of the IDR region and that two of our protein-coding isoforms contained only the TBP-binding region and IDR domains. IDRs in transcription factors drive the formation of liquid-liquid phase separation necessary for transcriptional machinery assembly^55,56^ and a disease associated repeat expansion in the IDR of the *Taf1* binding partner TBP likely exerts its pathogenic mechanism through changing phase separation capacity^57^. IDRs have also been suggested to be crucial for determining transcription factor binding specificity^58^. Therefore, given the observed protein diversity in this region and neuronal specificity of a microexon in the *Taf1* IDR, future work into how different *Taf1* IDRs influence its transcriptional function and phase-separation capacity will be critical to understanding both its normal and pathogenic roles.

To understand the function of individual genes it is essential to catalogue their RNA and protein outputs, spatially and temporally, ideally from each cell type, in healthy tissue. In addition, patterns of association between different exons and promoters can give new insight into gene function. AS can have a profound effect on gene/protein function for example, by altering stability, localisation and interactions^59,60^. Much remains to be experimentally determined about TAF1, a large and fundamentally important protein, including the roles of its multiple mRNA variants and protein isoforms.

## Materials and methods

### Mouse strain, welfare and husbandry

Female and male C57BL/6J wild-type mice (from The Jackson Laboratory) were maintained in Specific Pathogen-Free (SPF) conditions, in individually ventilated cages; mice were kept in adherence to the Home Office Code of Practice and licensed by the Home Office under the Animals (Scientific Procedures) Act 1986 Amendment Regulations 2012 (SI 4 2012/3039), UK. All experiments had Institutional Ethical Review Committee approval. Mice had access to a mouse house with bedding material, wood chips, and continual access to water and RM1 (Special Diet Services) (stock animals) chow. Animals were euthanized at 3-months, 6-months and 12-months of age by exposure to rising carbon dioxide, followed by confirmation of death by dislocation of the neck, in accordance with the Animals (Scientific Procedures) Act 1986 (UK).

### Total RNA extraction and First-Strand cDNA synthesis

Total RNA was extracted from 3-month-old mouse brain from four brain regions: cerebellum, cortex, hippocampus and striatum (Supplementary Table S1) using the RNeasy Mini Kit (Qiagen). Total RNA was extracted from 3-month-old mouse spleen, heart and quadricep muscle (Supplementary Table S1) using the RNeasy Fibrous Tissue Mini Kit (Qiagen); all tissue was homogenised on ice using a TissueRuptor II (QIAGEN) in Buffer RLT with 1% β-mercaptoethanol, following the manufacturer’s instructions for both kits. The final extracted RNA was eluted in DNase- and RNase-free water. RNA was quantified using a NanoDrop spectrophotometer followed by RNA Integrity Analysis. cDNA synthesis was carried out using Superscript™ IV First-Strand Synthesis System (11904-018; Thermo Fisher Scientific). 2 μg RNA was used as input for annealing of oligo-dT primers. Final clean-up was performed with 1 μL of RNase H.

### RNA integrity analysis

A volume of 1μL of undiluted RNA from each sample was run on the Agilent 4200 TapeStation using RNA screen tape (Agilent, Santa Clara, CA, USA cat #5067-5576). The concentrations of the samples ranged from 92.1 ng/μL to 1368.0 ng/μL, samples of >500 ng/μL were diluted to fall within the quantitative range for the assay (25–500 ng/μL). RINe was recorded from the TapeStation 4200 and assigned to the sample automatically. A cut-off value of RINe 7.5 determined whether samples could then be used.

### Long-range PCR amplification of mouse Taf1 cDNA

PCR was performed using LongAmp® *Taq* DNA Polymerase (New England Biolabs). Optimal annealing temperature was determined using a gradient thermocycler. Primers were designed and screened for specificity using NCBI’s Primer-BLAST. Primers annealled to non-coding regions (5′ and 3′ UTRs) flanking the longest common sequence of c*Taf1* (GenBank (NM_001045959.2). 5′ UTR Forward primer sequence: AAGTCACCCGTGTGCGACTGACG, through to 3′ UTR Reverse primer sequence: AGCTGGGGGGAAAGGTAATAGACC. For each sample, duplicate PCRs were performed in 50 μL volumes with 0.1 mM of each primer. Thermocycling conditions were: initial denaturation step at 94 °C for 30 seconds, followed by 29 cycles of 94 °C for 30 seconds for denaturation, 61 °C for 30 seconds for annealing, and 65 °C for 12 min for extension. This was followed by a final extension at 65 °C for 10 min, and reactions were held at 4 °C indefinitely. The number of cycles was optimized at 29 to ensure adequate amplification of the target sequences while minimizing non-specific products. Following amplification, duplicate samples were pooled and the size of the RT-PCR product verified by agarose gel electrophoresis in 1x TAE buffer using SYBR Safe DNA stain for visualisation. The remaining PCR product (∼80 μL) was sent to Full Circle Laboratories and purified using 0.5x AMPure XP beads (Beckman Coulter, Fullerton, USA) before proceeding to library construction.

### Library preparation and nanopore sequencing

Nanopore sequencing libraries were prepared and barcoded using the Native Barcoding Kit 96 V14 (SQK-NBD114.96). 4.15 μL of purified amplicon DNA was subjected to an end-prep reaction using the NEBNext Ultra II End Repair/dA-tailing Module (New England BioLabs, MA). End-prepped amplicons were taken forward to the native barcode ligation step; a total of 0.75 μL of end-prepped amplicons were mixed with 1.25 μL of native barcode, 5 μL of blunt/TA ligase master mix, and 3 μL H_2_O and then incubated at room temperature for 20 min. After incubation 2 μL of 0.25 M EDTA was added to each sample to stop the ligation reaction. The barcoded amplicons were pooled, and a 0.5x AMPure XP bead (Beckman Coulter, CA) purification was carried out with two subsequent 500 μL 80% ethanol washes. The barcoded library was eluted in 30 μL nuclease-free water, then subjected to ligation of the sequencing adapter; 10 μL of 5× NEBNext quick ligation reaction buffer, 5 μL ONT *Adapter Mix* II (*AMII*), and 5 μL Quick T4 DNA ligase were mixed with 30 μL of the barcoded library and then incubated at room temperature for 20 min. The library was purified using 0.4× AMPure XP beads with two subsequent 125 μL Long Fragment Buffer (LFB) washes and eluted in 30 μL EB buffer (ONT). The final library was quantified with the Qubit 1× dsDNA high-sensitivity assay kit (Invitrogen) and a Qubit 4 fluorometer (Invitrogen). Approximately 60 ng of the final library was loaded onto the FLO-PRO114M (R10.4.1) flow cell on an ONT PromethION 2 Solo platform for 10 h.

### Long-read cDNA-seq bioinformatics: alignment and microexon refinement

Raw ONT sequencing reads in BAM files were aligned to the *Mus musculus* GRCm39 primary assembly reference genome using minimap2 (v2.17) with the spliced alignment preset, optimized for long-read transcriptomic data. To improve alignment accuracy for microexons, particularly *Taf1* microexon 34′, alignments were refined using MisER. Resulting alignments in SAM format were converted to BAM and indexed by samtools (v1.15). For targeted inspection of the *Taf1* locus, alignments were filtered for reads mapping specifically to chromosome X coordinates chrX:100576335-100644635 (GRCm39). Manual inspection of exon-intron architectures, primer binding sites, and candidate novel exons was conducted using the Integrative Genomics Viewer (IGV v2.18.4). Summary of sequence data: Supplementary Table 2.

### Long-read cDNA-seq bioinformatics: transcript identification and quantification

Transcript identification was performed using IsoQuant (v3.2.0), optimized for transcript-level analysis of long-read sequencing data. IsoQuant was executed with the ONT sequencing data flag and configured to generate SQANTI-compatible outputs. Novel transcripts identified by IsoQuant were organized in sample-specific directories and merged across samples using StringTie2. The merged annotation in GTF format was converted to FASTA format using gffread for quantification. Long-read transcript quantification was performed using Oarfish (v0.9.0) with model coverage applied and no filters, employing the merged transcript reference FASTA file. All computational analyses were conducted within a dedicated, reproducible Conda environment.

### Transcript expression analysis

Differential transcript usage and differential transcript expression were conducted using the R packages tximeta and Swish method (with fishpond v2.14.0). Transcript quantification files from Oarfish were imported into R using tximeta. For DTU and DTE, Swish, using scaling based on library size and effective transcript length, yielded log2-scaled counts for nonparametric differential analysis.

### Protein isolation and western blotting of mouse brain

For protein isolation, one brain hemisphere was dissected in ice-cold PBS to isolate cerebellum, cortex, hippocampus and striatum (Supplementary Table S1). All tissues were then snap frozen in liquid nitrogen. Samples were homogenised in ice-cold RIPA Lysis Buffer (Millipore) plus Protease Inhibitor Cocktail I (Millipore, MA, USA) by mechanical disruption using a TissueRuptor II (QIAGEN). Homogenates were shaken on ice at 70 rpm for 2 h at 4 °C, then centrifuged for 20 min at 15,000 *g* at 4°C. Supernatants were removed and stored at −80 °C until use. Total protein concentration was determined by Bradford assay using Protein Assay Dye Reagent (Bio-Rad, CA, USA). Equal amounts of total brain protein were denatured in LDS denaturing buffer (Thermo Fisher Scientific) and β-mercaptoethanol and incubated at 95°C for 5 min prior to separation by SDS-PAGE gel electrophoresis.

Protein samples (40 μg) were loaded onto precast NuPAGE 7% Tris-Acetate gels (Thermo Fisher Scientific) and electrophoresed at 80 V for 4 h in an ice bath using NuPAGE Tris-Acetate Running Buffer (Thermo Fisher Scientific). Proteins were transferred to 0.2 μm nitrocellulose membranes using a Trans-Blot Turbo transfer system (Bio-Rad) High Molecular Weight Program at 1.3 A for 30 min. Membranes were stained for total protein with 0.1 % (w/v) Ponceau S in 1 % (v/v) acetic acid for 5 min prior to blocking for 1 h at room temperature in Intercept PBS Blocking Buffer (LI-COR Biosciences, NE, USA). Membranes were incubated overnight in primary antibodies diluted in Intercept PBS Blocking Buffer (LI-COR Biosciences) at 4 °C (TAF1 [177-4], 1:1000; β-tubulin, 1:10,000). Membranes were then washed three times for 10 min in PBS containing 0.05 % Tween-20 (PBS-T) and incubated for 1 h at room temperature with the relevant IRDye secondary antibody (LI-COR Biosciences) diluted in Intercept PBS Blocking Buffer. Finally, membranes were washed three times for 10 min in PBS-T and once for 10 min in PBS. Antibody binding was visualised using the Odyssey CLx Imaging System (LI-COR Biosciences) and ImageJ was used for band density analysis and total protein quantification. All antibodies used are described in Supplementary Table S3.

Reproducible results were obtained through multiple blots, and the experiment was repeated for samples that showed large variation in protein expression, which, following quantification, were excluded as outliers in the Grubb’s outliers test, but no samples were excluded from the analysis. No randomisation was used, and the investigator was aware of the genotype.

Relative signal of the TAF1 antibody compared to an internal loading control (β-tubulin) was calculated and the values were normalised to the mean relative signal of control samples electrophoresed on the same gel. The mean of technical replicates was calculated and used for ANOVA, for which biological replicates were used as the experimental unit and the assumptions for this test were met.

### Protein domain prediction

To characterise novel transcripts identified in the Oarfish transcriptome, protein structural analyses were conducted utilizing the IsoformSwitchAnalyzeR package (v2.8.0) in R using a custom GTF file, integrating novel Isoquant transcripts and known *Taf1* transcripts. ORFs were predicted with reference to the mouse genome (BSgenome.Mmusculus.UCSC.mm39, Gencode). The “mostUpstream” parameter was employed to select ORFs, ensuring identification of the earliest start codon and minimizing the incorporation of downstream sequences potentially encoding unrelated proteins.

Transcripts were subsequently evaluated for potential NMD through detection of predicted PTCs. Amino acid sequences derived from predicted-ORFs were extracted via the “extractSequence()” function and subjected to domain annotation. Protein domains were identified using PFAM scan (v1.6) with default parameters. Intrinsically disordered regions (IDRs) were predicted using IUPred2A (https://iupred2a.elte.hu/). Signal peptide sequences, indicating potential subcellular localization or secretion, were predicted using SignalP (v6.0).

### TAF1 protein complex modelling

The complex between TAF1 (Uniprot Q80UV9), TBP (P29037), TAF11 (Q99JX1) and TAF13 (P61216) was modelled using the Alphafold 3 server^61^ and visualized in Pymol 3.1.6. Complexes containing the truncated TAF1.1 and TAF1.14 splicing transcripts produced the same structural model with shortened loop regions.

### Immunohistochemistry

Three male and three female mice, 3-months of age, were euthanised and perfused with PBS to ensure complete removal of blood. This was followed by perfusion with 50 ml 4% paraformaldehyde. Brains were extracted and fixed in 4% paraformaldehyde for 24 h at 4°C. Post-fixation, samples were washed twice in PBS and stored in PBS containing 0.05% sodium azide at 4°C. Brains were sagittally sectioned using a vibrating microtome (Leica VT1000S) at a thickness of 40 μm and maintained in PBS with 0.05% sodium azide at 4°C until staining. Immunohistochemistry was conducted following the protocol outlined by Crombie et al.^1^ Sections were blocked and permeabilized in a solution of 5% goat serum with 0.25% Triton X-100 for 1 h. Sections were then incubated with primary antibodies (TAF1 [177-4] at 1:200; DARPP-32 [H-3] at 1:1000) in 5% goat serum overnight at 4°C. Negative control sections were treated with 5% goat serum alone. Following primary antibody incubation, all samples were washed three times for 5 min each in PBS. Sections were then incubated with secondary antibodies (goat anti-rabbit IgG (H+L) Alexa Fluor 564 at 1:400; goat anti-mouse IgG (H+L) Alexa Fluor 488 at 1:1000) for 1 h. After secondary antibody incubation, samples were washed three times in PBS and incubated in DAPI solution for 5 min. Finally, samples were mounted using Prolong Gold Antifade (P10144, Vector Laboratories) and coverslipped. Imaging was conducted at high magnification using a Zeiss LSM980 confocal microscope, with whole sections imaged utilizing an Axio Scan Z1 slide scanner.

### Statistical analysis

To compare *Taf1* expression across the four different brain regions, an ordinary one-way ANOVA was conducted. For DTU and DTE analyses, Swish (with fishpond) performed nonparametric testing using the Wilcoxon rank-sum test for pairwise comparisons between conditions. Statistical significance was defined as p < 0.05 (*p-value < 0.05; **p-value < 0.01; ***p-value < 0.005). Western blot quantification and statistical analyses were performed using GraphPad Prism 10, accessed in August 2024.

## Data and Code Availability

RNA sequencing data are available through the ArrayExpress under accession E-MTAB-15282. The codes used in this study are deposited in https://github.com/PeihangLi/Taf1-longreads. No new resources were generated in this study.

## Authors’ contributions

PL, KC, JB and A-LB undertook data curation and formal analysis. All authors contributed to the conceptualisation and investigation. A-LB and EMCF undertook funding acquisition. All authors contributed to all stages of writing and have read and approved the final manuscript.

## Acknowledgments

EMCF, PL and KC were supported by the Collaborative Center for X-linked Dystonia–Parkinsonism (no. 239295 to EMCF). JB was supported by the Medical Research Council (MC_UU_00037/7) and the National Institute of Neurological Disorders and Stroke (1R21NS101588-01A1). A-LB was supported by a Fellowship from the Guarantors of Brain.

We thank Marc Timmers for useful discussions, and for the TAF1 [177-4] antibody, produced by the Timmers’ laboratory with support from the Collaborative Center for X-linked Dystonia-Parkinsonism (CCXDP).

---

**Supplementary Table S1.** C57BL/6J Mouse Biological Samples.

| Sex | ID | Age (m) | Tissue |
| --- | --- | --- | --- |
| Male | 913001 | 3 | Striatum |
|  |  |  | Cerebellum |
|  |  |  | Hippocampus |
|  |  |  | Cortex |
|  | 913008 | 3 | Heart |
|  |  |  | Muscle |
|  |  |  | Spleen |
|  | 913004 | 3 | Striatum |
|  |  |  | Cerebellum |
|  |  |  | Hippocampus |
|  |  |  | Cortex |
|  | 913009 | 3 | Heart |
|  |  |  | Muscle |
|  |  |  | Spleen |
| Female | 912979 | 3 | Striatum |
|  |  |  | Cerebellum |
|  |  |  | Hippocampus |
|  |  |  | Cortex |
|  | 912986 | 3 | Heart |
|  |  |  | Muscle |
|  |  |  | Spleen |
|  | 912985 | 3 | Striatum |
|  |  |  | Cerebellum |
|  |  |  | Hippocampus |
|  |  |  | Cortex |
|  | 912987 | 3 | Heart |
|  |  |  | Muscle |
|  |  |  | Spleen |

**Supplementary Table S2.**
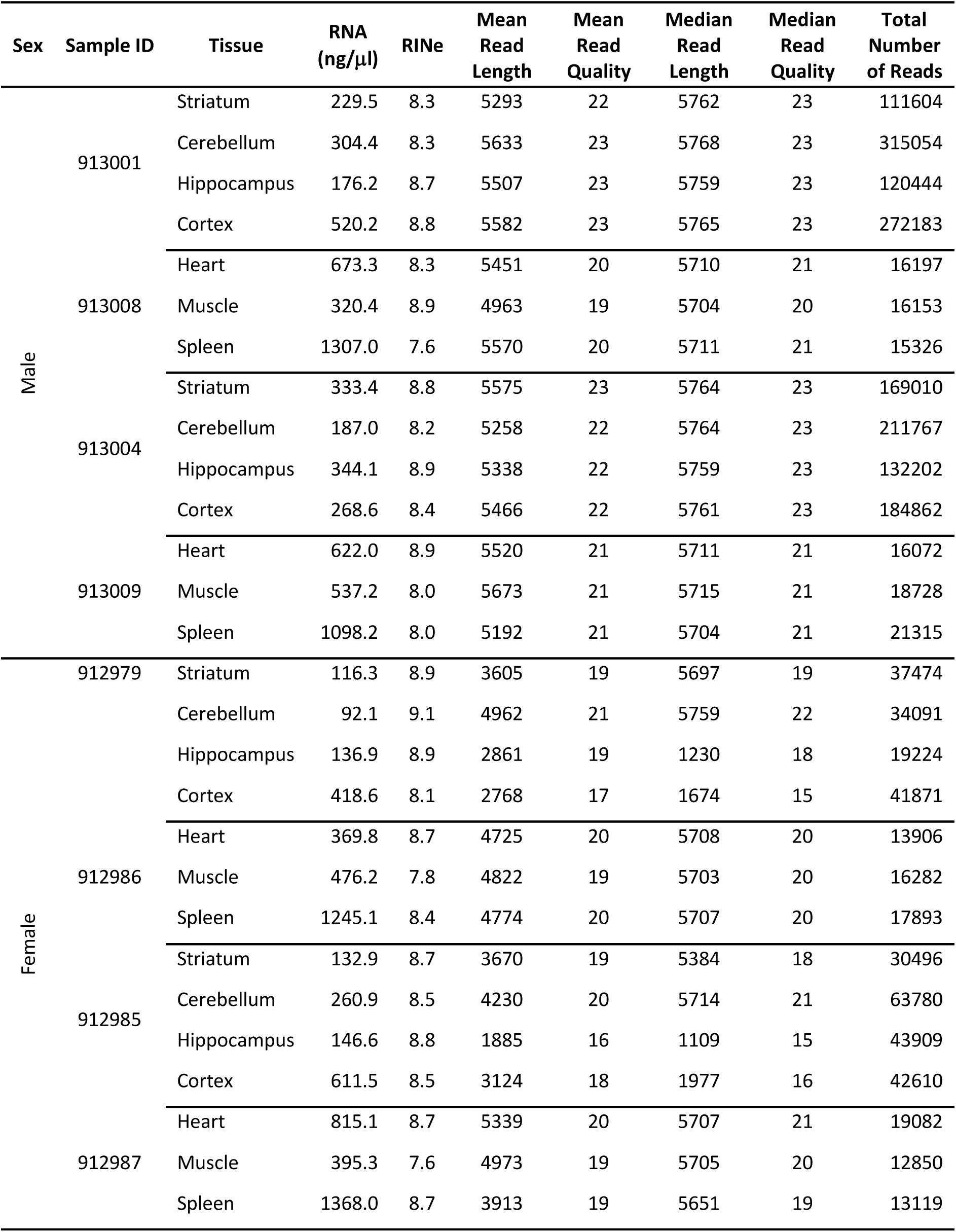
Summary statistics of sequencing.

**Supplementary Table S3.** Antibodies. Antibodies used for immunofluorescent staining (IF) and western blots (WB). Dilutions are described in Materials and Methods

| Category | Target (clone) | Company | Catalogue no. | Batch no. | Application |
| --- | --- | --- | --- | --- | --- |
| Primary | TAF1 [177-4]*<br>(rabbit monoclonal) | Timmers Lab<br>supported by<br>CCXDP (Capponi<br>et al. (2020)) | N/A (not<br>commercially<br>available) | N/A | WB/IF |
| Primary | Primary β-tubulin<br>[EPR16774]<br>(rabbit monoclonal) | Abcam | ab179513 | GR3214136-5 | WB |
| Secondary | Anti-rabbit IgG (H+L)<br>IRDye 800CW | LI-COR | 926-32211 | D10512-05 | WB |
| Secondary | Anti-mouse IgG<br>(H+L) IRDye 680LT | LI-COR | 926-68020 | D11005-15 | WB |
| Primary | CTIP2 [25B6]<br>(rat monoclonal) | Abcam | Ab18465 | 2101032938 | IF |
| Primary | DARPP-32 [H-3]<br>(mouse monoclonal) | Santa Cruz<br>Biotechnology | sc-271111 | 1021 | IF |
| Secondary | Anti-rat IgG (H+L)<br>Alexa Fluor 488 | Thermo Fisher | A-11006 | 2551392 | IF |
| Secondary | Anti-mouse IgG<br>(H+L) Alexa Fluor 488 | Thermo Fisher | A-11001 | 2551357 | IF |
| Secondary | Anti-rabbit IgG (H+L)<br>Alexa Fluor 564 | Thermo Fisher | A-11035 | 2387450 | IF |

\* Validation of this TAF1-specific monoclonal antibody has been published in Capponi et al. (2020). The epitope is TPGPYTPQPPDLY derived from peptides encoded by exons 34 and 35 of *TAF1/Taf1* (*Taf1*-201 and *Taf1*-202)

**Supplementary Table S4.** Coding region similarity: novel vs c*Taf1* (%)

| Coding region similarity: novel vs *cTaf1* (%) |  |
| --- | --- |
| Novel *Taf1* transcripts | Mouse *cTaf1* |
| *MSTRG.1.1* | 97.73 |
| *MSTRG.1.3* | 97.04 |
| *MSTRG.1.7* | 97.14 |
| *MSTRG.1.8* | 96.05 |
| *MSTRG.1.11* | 69.53 |
| *MSTRG.1.14* | 97.46 |
| *MSTRG.1.17* | 30.34 |
| *MSTRG.1.19* | 14.60 |

**Supplementary Table S5:** Length of novel coding exons.

| Novel coding exons | Length (bp) |
| --- | --- |
| truncated exon 4 | 42 |
| truncated exon 5 | 179 |
| full-length exon 34'' | 172 |
| truncated exon 34'' | 105 |

**Figure S1:**
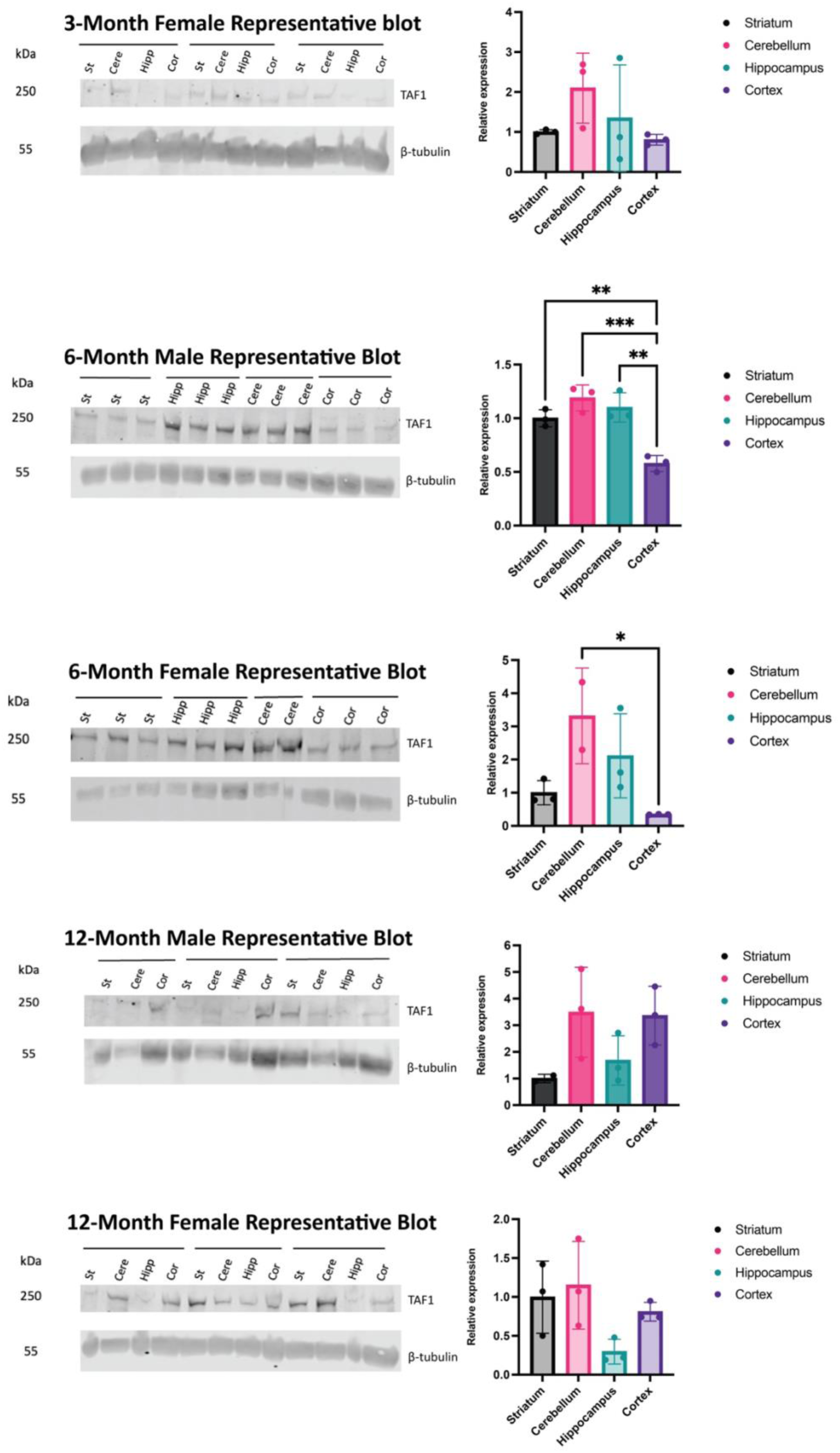
TAF1 protein expression in mouse brain. Comparison of TAF1 protein expression in striatum, cerebellum, hippocampus and cortex in littermates at 3-months, 6-months and 12-months of age, *n*=3/brain region/sex). Error bars show mean ± standard deviation of the relative expression levels. Statistical analysis using one-way ANOVA was performed; *p*-values are indicated. The experiment was replicated three times or more. For each age and sex, representative blots are shown on the left, with a β-tubulin loading control. Data for each brain region are color-coded; striatum (black), cerebellum (pink), hippocampus (blue), cortex (purple). Significant differences between brain regions within the same sex and age group are indicated by asterisks, with statistical significance denoted as \**p* < 0.05, \*\**p* < 0.01, and \*\*\**p* < 0.001.

**Figure S2.**
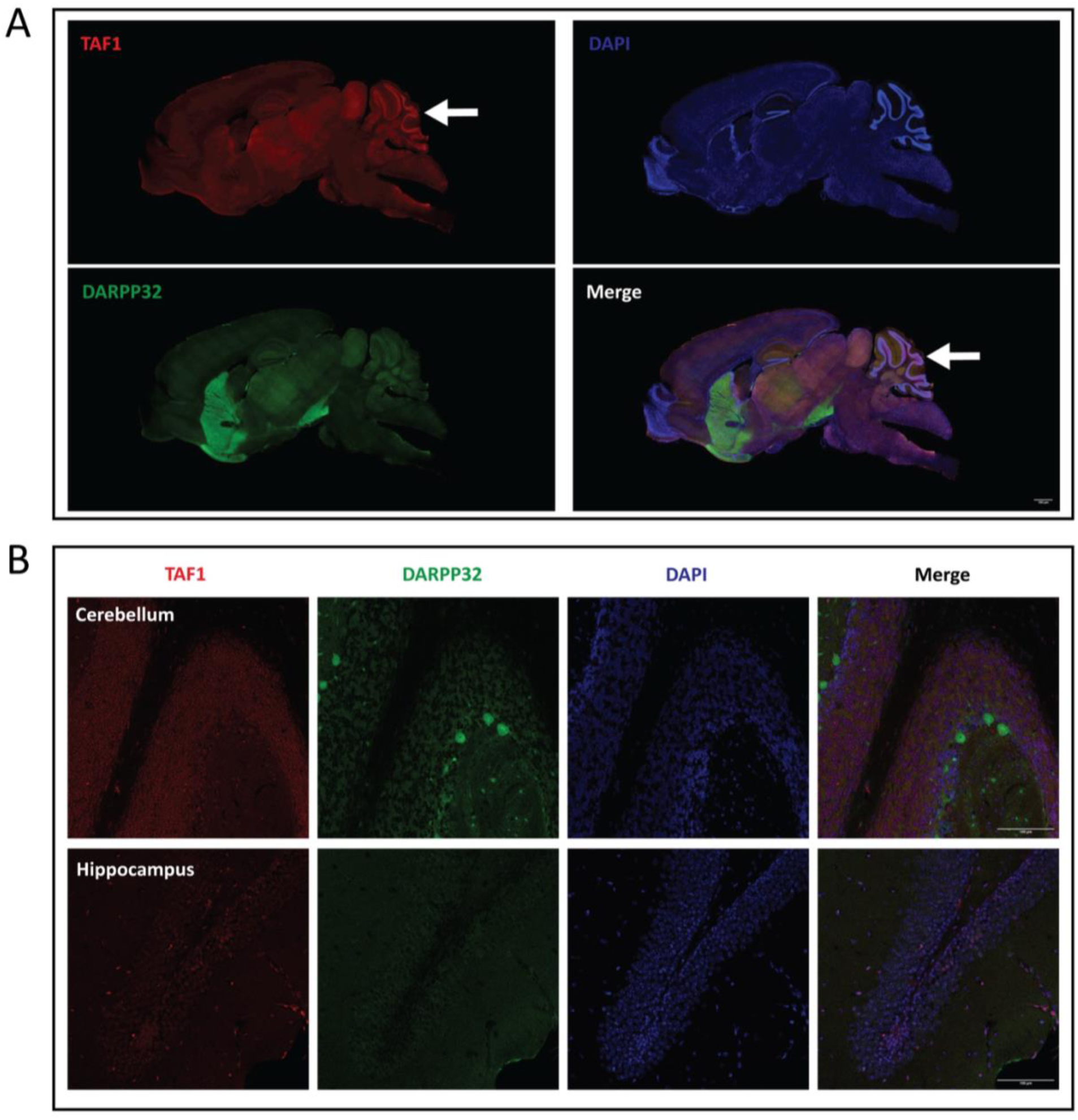
TAF1 expression in female mouse brain. Immunofluorescent staining of 3-month-old female C57BL/6J mice. (A) Sagittal brain sections, (B) sections of cerebellum and hippocampus. Sections are stained for TAF1 (red), DARPP-32 (green), and DAPI (blue), with the merged images showing the colocalization of these markers. White arrow indicates cerebellum. The experiment was repeated twice with reproducible results. This was not a quantitative experiment.

